# Proteomic analysis revealed the function of PoElp3 in development, pathogenicity, and autophagy through the tRNA-mediated translation efficiency in the rice blast fungus

**DOI:** 10.1101/2023.07.19.548190

**Authors:** Yuanhao Liu, Ting Sun, Yuyong Li, Jianqiang Huang, Xianjun Wang, Huimin Bai, Jiayi Hu, Zifan Zhang, Shuai Wang, Dongmei Zhang, Xiuxiu Li, Zonghua Wang, Huakun Zheng, Guifang Lin

## Abstract

The Elongator complex is conserved in a wide range of species and plays crucial roles in diverse cellular processes. We have previously shown that the Elongator protein PoELp3 was involved in the asexual development, pathogenicity, and autophagy of the rice blast fungus. In this study, we further revealed that PoElp3 functions via tRNA-mediated protein integrity. Phenotypic analyses revealed that overexpression of two of the tRNAs, tK(UUU) and tQ(UUG) could rescue the defects in Δ*Poelp3* strain. TMT-based proteomic and transcriptional analyses demonstrated that 386 proteins were down-regulated in Δ*Poelp3* strain compared with wild type strain Guy11, in a transcription-independent manner. Codon usage assays revealed an enrichment of Glutamine CAA-biased mRNA in the 386 proteins compared with the 70-15 genome. In addition to those reported previously, we also found that PoErp9, a sphingolipid C9-methyltransferase, was down-regulated in the Δ*Poelp3* strain. Through an *ILV2*-specific integration of *PoERP9-GFP* into the wild type and Δ*Poelp3* strain, we were able to show that PoErp9 was positively regulated by PoElp3 translationally but not transcriptionally. Functional analyses revealed that PoErp9 was involved in the fungal growth, conidial development, pathogenicity, and TOR-related autophagy homeostasis in *P. oryzae*. Taken together, our results suggested that PoElp3 acts through the tRNA-mediated translational efficiency to regulate asexual development, pathogenicity, and autophagy in the rice blast fungus.

## 1. Introduction

Transfer RNAs (tRNAs) are central players in protein biosynthesis that decipher the genetic code of mRNA and add appropriate amino acids into the growing amino acid chain (Holley *et al*. 1965; Ma *et al*. 2021). The simple primary sequence of tRNAs determines that it could fold into a diverse structure in addition to the one with biological function. Therefore, the posttranscriptional modifications of tRNA are essential for the maturing, folding, degradation, and decoding efficiency (Sokołowski *et al*. 2018). The 5-methoxycarbonylmethyl-2-thiouridine (mcm^5^s^2^) modification of tRNA at U_34_ is one of the best-studied modifications and is conserved in many species (Leidel *et al*. 2009; Sokołowski *et al*. 2018). The mcm^5^s^2^-modified tRNAs decipher the NAA type codons and wobble onto NAG. For example, the lysine, glutamine, and glutamic acid are encoded by either AAA, CAA, and GAA, or AAG, CAG, and GAG, respective. And a single eukaryotic tRNA (tRNA^Lys^ _UUU_ (tK^UUU^), tRNA^Glu^ _UUC_ (tE^UUC^), or tRNA^Gln^ _UUG_ (tQ^UUG^)) with a U at position 34 can recognize both the NAA and NAG (wobble) codons (Dewez *et al*. 2008; Goffena *et al*. 2018). The Elongator complex is involved in the first step of mcm^5^s^2^ modification by transferring a carboxymethyl group to U_34_ at the C_5_ position to form the 5-carboxymethyluridine (cm^5^U) precursor (Selvadurai *et al*. 2014). Although Elongator complex was found to modify 11 U_34_-carrying tRNA species, only tK^UUU^, tE^UUC^, and tQ^UUG^ were subsequently thiolated (S2) (Johansson *et al*. 2008).

Elp3 is the catalytic subunit of the Elongator complex, which was originally identified in *Saccharomyces cerevisiae* as a component of elongating RNA polymerase II (RNAPII) holoenzyme (Otero *et al*. 1999). Elp3 contains a Gcn5-related N-terminal histone acetyltransferase (HAT) domain (Winkler *et al*. 2002), which was reported to have the acetyltransferase activity on histone protein and other proteins (Creppe *et al*. 2009; Miśkiewicz *et al*. 2011; Laguesse *et al*. 2017; Lin *et al*. 2019; Planelles-Herrero *et al*. 2022). However, the effect of Elp3 on histone acetylation could be altered in different organisms, in suggestive of diverse regulatory mechanisms in different backgrounds. For example, the H3K14ac level was significantly reduced in *S. cerevisiae* (Wittschieben *et al*. 2000), *Fusarium graminearum* (Lee *et al*. 2014) and *Beauveria bassiana* (Cai *et al*. 2022), but not in *Aspergillus fumigatus* (Zhang *et al*. 2022) and *Pyricularia oryzae* (Zhang *et al*. 2021). Elp3 also possesses a radical S-adenosylmethionine (SAM) domain, which is responsible for the modification of wobble uridine residues in eukaryotic tRNA (Dauden *et al*. 2019; Lin *et al*. 2019; Abbassi *et al*. 2020). Increasing evidence demonstrated that overexpression of corresponding hypomodified tRNA could rescue the defects caused by Elongator complex-associated s^2^U_34_ deficiency in many organisms, such as *S. cerevisiae* (Esberg *et al*. 2006; Nedialkova and Leidel 2015; Tükenmez *et al*. 2015; Jaciuk *et al*. 2023), *Caenorhabditis elegans* (Nedialkova and Leidel 2015), and *Aspergillus fumigatus* (Zhang *et al*. 2022). In line with this, studies in different species demonstrated that both the SAM and KAT domains are required for the modification of tRNA by Elp3 proteins (Selvadurai *et al*. 2014; Zhang *et al*. 2022). These results further support a key role of Elongator complex in maintaining the wobbling function of tRNA.

In *P. oryzae* (syn. *Magnaporthe orzyae*), causal agent of the destructive blast disease, autophagy is crucial for diverse developmental processes, particularly, the appressorial development (Liu *et al*. 2007; Wilson and Talbot 2009; He *et al*. 2018; Sun *et al*. 2018; Yin *et al*. 2019). We had previously shown that the Elp3 homolog was required for the development and pathogenicity in the rice blast fungus *P. oryzae*, probably by maintaining the autophagy homeostasis (Zhang *et al*. 2021). In this study, we aim to investigate whether PoElp3 is involved in the wobbling function of tRNA and subsequently the proteomic integrity.

## 2. Materials and methods

### 2.1. Manipulation of fungal strains

The fungal strains were cultivated as described previously (Anjago *et al*. 2022). In brief, all strains were grown at 28 °C on complete medium (CM, containing yeast extract 0.6%, casamino acid 0.6%, sucrose 1%, agar 2%). For rapamycin inhibition assay, the strains were grown at 28 °C on MM medium supplemented with or without 1 μg/mL rapamycin (He *et al*. 2018). All the assays were repeated three times with at least three replicates in each biological repeat.

### 2.2. Spray inoculation assay

The spray inoculation assays were performed as previously described with minor modification (Zhang *et al*. 2021). Conidial suspensions (5×10^4^ conidia/mL in 0.025% Tween-20 solution) of each strain were used for spray inoculation assay on 14-day-old TP309 (Fig. 1) or NPB (Fig. 6) rice seedlings. The challenged rice seedlings were then cultivated under the conditions described previously (Giraldo *et al*. 2013). The symptoms were recorded and scored as previously described (Valent *et al*. 1991).

**Fig. 1.**
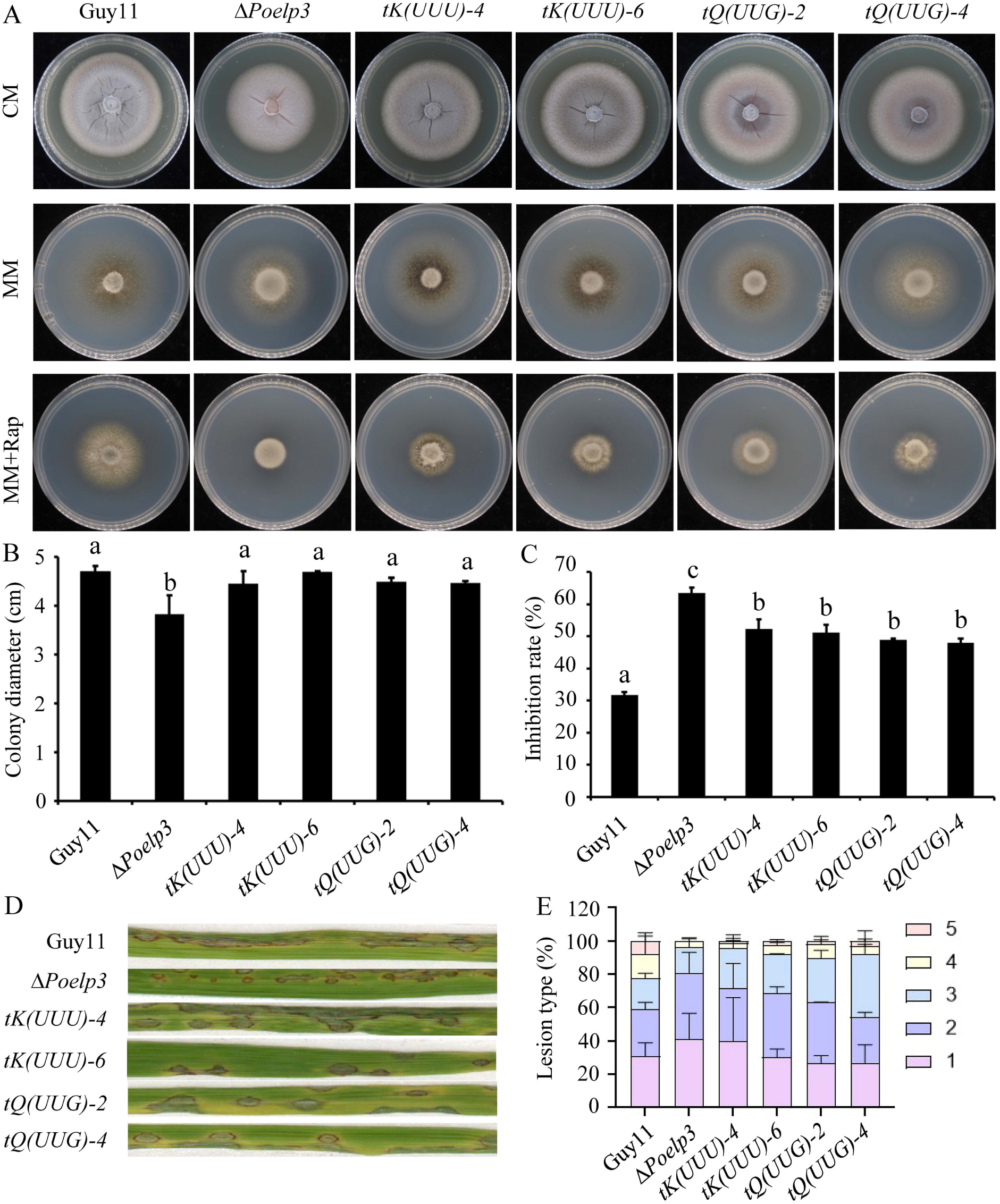
Overexpression of *tK*(*UUU*) or *tQ*(*UUG*) could partially rescue the defects in Δ*Poelp3* strain. A, the indicated strains grown on CM, MM and MM supplemented with 1 μg/uL rapamycin (MM+Rap). B, colony diameter of indicated strains grown on CM. C, the inhibition rate of indicated strains upon the treatment of 1 μg/uL rapamycin. The letters indicate statistically significant differences (p<0.01). D, leaves of rice cultivar TP309 were sprayed with conidial suspensions (5×10^4^conidia mL^−1^) of strains indicated. E, quantification of different lesion types. Data were shown as the mean±SE (n=3).

### 2.3. Construction of tRNA overexpression strains

For the identification of tRNA^Lys^ _UUU_ (tK^UUU^) and tRNA^Gln^ _UUG_ (tQ^UUG^), we search the rice blast genome for the homologs of SPBTRNALYS.06 and SPBTRNAGLN.02, respectively, from *Schizosaccharomyces pombe* (Appendix A). The identified coding sequences of each tRNA were amplified using the primers listed in Appendix B and inserted into the *Xho*I and *Hind*III sites of pKNT-RP27, which was then introduced into the Δ*Poelp3* strain through the PEG3350-meidiated protoplast transformation (Sweigard *et al*. 1998). The resulting transformants were selected on TB3 medium with 600 μg/mL G418 and further tested through PCR-based genotyping.

### 2.4. Proteomics analysis

The proteomic analysis was performed as previously described with some modifications (Zhang *et al*. 2022). Tandem Mass Tags (TMT)-based proteomics analysis was conducted by Jingjie PTM Bio (Hangzhou, China). 50 μg total protein of each sample was digested with trypsin for TMT labeling and fraction. The LC-MS/MS analysis was performed using an Orbitrap Exploris™ 480 mass spectrometer (Thermo Scientific™, USA). The MS raw data were processed with Proteome Discoverer (v2.4.1.15) and searched against 70-15 reference genome (Dean *et al*. 2005).

### 2.5. RNA-sequencing and data analysis

The RNA-sequencing and data analysis were performed as described previously (Xie *et al*. 2023). The total RNA of each sample was used for the construction of the RNA-seq library as described previously (Zhong *et al*. 2019). The libraries were sequenced using the Illumina NovaSeq 6000 system. Quality assessment of raw reads (RNA-seq) was conducted using FASTQC v0.11.3 (www.bioinformatics. babraham.ac.uk/bugzilla/). Adapters were removed from the raw reads using Trim_galore v3.1 (https://github.com/FelixKrueger/TrimGalore/). Then, the RNA-seq reads were aligned to 70-15 reference genome (Dean *et al*. 2005) using RSEM v1.3.3 (Li and Dewey 2011). The quantification of expression counts were conducted with transcript per million (TPM).

### 2.6 Generation of the *ILV2*-specific *PoERP9-GFP* integration strains

The construction of the *ILV2*-specific *PoERP9-GFP* integration vector and the generation of *ILV2*-specific *PoERP9-GFP* integration strains was performed as described previously (Yang and Naqvi 2014). The native promoter and coding sequence were amplified respective using the primers listed in appendix B and inserted into *pKNT-GFP*. The resultant construct *pKNT-PoERP9p-PoERP9-GFP* was used as the template for the amplification of *PoERP9p-PoERP9-GFP* fusion, which was them integrated into pFGL1010-GFP. The resultant construct pFGL1010-PoERP9p-PoERP9-GFP was transformed into the *Agrobacterium tumefaciens* AGL1 strain via electroporation. The *A. tumefaciens*-mediated transformation (ATMT) of rice blast fungus was performed as described previously (Yang and Naqvi 2014).

### 2.7. Targeted gene deletion and complementation of *PoERP9*

Strategy for the targeted gene deletion of *PoERP9* and the construction of complemented strain (Δ*Poerp9c*) was performed as described previously (Zhang *et al*. 2021). In brief, the upstream fragment (A) and downstream fragment (B) were amplified using *ERP9AF* & *ERP9AR*, and *ERP9BF* & *ERP9BR* (Appendix B), which were then fused with the 5’-terminal (HY) and 3’-terminal (YG) of hygromycin B phosphotransferase cassette (HPH), respectively, to form the ‘A-H’ and ‘H-B’ fusion constructs (Appendix C), respectively. The ‘A-H’ and ‘H-B’ fusions were introduced into the *Ku80* protoplasts using the PEG-mediated transformation method as described previously (Kershaw and Talbot 2009). PCR-based genotyping of the deletion strain was performed using the primer pairs *ERP9tF* & *ERP9tR*, and *ERP9UAF* & *H853* (Appendix B). Southern blot assay was performed for to validate the mutant candidates.

For complementation of Δ*Poerp9* strain, genomic DNA fragment containing the native promoter, ORF region, and 3’-untranslated region (UTR) of *PoERP9* was amplified from the Guy11 genomic DNA using primers *ERP9comF* and *ERP9comR* (Appendix B) and inserted into *pKNTG* vector carrying the neomycin resistance cassette. The resulting construct was transformed into the Δ*Poerp9* protoplasts as described above. The neomycin resistant colonies were evaluated through PCR-based identification using the primer pairs *ERP9tF* & *ERP9tR*.

### 2.8. Southern hybridization assay

10 μg genomic DNA isolated from the indicated strains was digested with *Kpn* I for the validation of Δ*Poerp9* and *Sac* I for the identification of *ILV2*-specific integration of *PoERP9-GFP*. The digested products were electrophoresed in 1% (w/v) agarose gel and then transferred onto positively charged Nylon membranes (Roche, Germany). For both assays, the ‘A’ fragment amplified from the Guy11 genomic DNA using primer pairs *ERP9AF*&*ERP9AR* was used as probe. Probe labeling, hybridization, and detection were performed using DIG High Prime DNA Labeling and Detection Starter Kit I (Roche) according to the instruction manual.

### 2.9. Western hybridization assay

The Western blotting assays were performed as described previously (Zhang *et al*. 2021). The mycelial pellets were ground into fine powder in liquid nitrogen and transferred into 1 mL extraction buffer (10 mmol L^−1^ Tris-HCl pH 7.5, 150 mmol L^−1^ NaCl, 0.5 mmol L^−1^ EDTA, 0.5% NP-40, 2 mmol L^−1^ PMSF). After the resuspension and centrifuge, SDS-PAGE was performed to resolve the proteins. For western blot assays, the resolved protein samples were transferred onto a PVDF membrane (Merck Millipore Ltd., USA) through a Bio-Rad electrophoretic blotting apparatus. The detection of PoErp9-GFP fusion and Actin was performed as described previously with minor revision (Zhang *et al*. 2021). The whole-cell extract detection assays were carried out with an anti-GFP antibody (Abmart, China), anti-Actin antibody (Huadingbio, China), and an anti-mouse secondary antibody (Abmart, China). After the incubation with the antibodies, the Western Bright ECL HRP substrate (Advansta, Menlo Park, USA) was used to detect the chemiluminescent signals.

### 2.10. Quantitative real-time PCR (qRT-PCR)

First strand cDNA was generated from 1 μg of total RNA using the Prime Script RT Reagent Kit with gDNA Eraser (TaKaRa, Dalian, China). Real-time PCR was performed by using an SYBR Green PCR Master Mix (TaKaRa, Dalian, China) and the primers listed in Appendix B.

### 2.11. Live-cell imaging assay

To image invasive hyphae, spore solution (10^5^ sporse mL^-1^ in ddH2O) of each strain was inoculated into the leaf sheath of susceptible rice cultivar CO39. For rapamycin inhibition assay, conidial suspension with or without 1 μg mL^−1^ rapamycin was inoculated onto the leaf sheath of CO39. The invasive hyphae were recorded at 24 hpi using Nikon A1R laser scanning confocal microscope system (Nikon, Japan) as previously reported (Zheng *et al*. 2017).

### 2.12. Statistical analysis

Statistical analysis was conducted by R packages in the R studio (Version 1.4.1106) environment (Xie *et al*. 2023).

## 3. Results

### 3.1. Overexpression of tRNA could rescue the defects in Δ*Poelp3* strain

To test whether the overexpression of tRNA could complement the defects showed in Δ*Poelp3* strain, we search in the rice blast fungal genome for genes encoding tK^UUU^ and tQ^UUG^, the two tRNA that could rescue the transcription and histone acetylation defects, as well as other phenotypes caused by the absence of Elongator when overexpressed (Esberg *et al*. 2006). We found that both tRNAs have two coding genes, with tK^UUU^-4 and tK^UUU^-6 localized at chr4 and chr6, tQ^UUG^-2 and tQ^UUG^-4 localized at chr2 and chr4, respectively. The coding sequence of the two copies of each tRNAs was highly conserved, while the franking sequences were highly polymorphic (Appendix A). The four tRNA coding genes were fused with the ribosomal protein promoter (*RP27p*) and introduced into the Δ*Poelp3* strain, respectively. At 7 days post inoculation, the diameter of transfromants carrying the *RP27p* :: *tK^UUU^* (herein after referred to as ‘*tK(UUU)*’) or *RP27p* :: *tQ^UUG^* (herein after referred to as ‘*tQ(UUG)*’) was about 95%-100% of that of wild type, while the diameter of Δ*Poelp3* strain was about 85% of that of the wild type (Fig. 1-A and B). Upon the treatment of rapamycin, inhibitor of the Target-of-Rapamycin (TOR) signaling pathway, the inhibition rate of fungal growth in the *tK^UUU^* and *tQ^UUG^* strains was reduced compared to Δ*Poelp3* strain, but was still larger than that of Guy11 (Fig. 1-A and C). Spray inoculation results showed that expression of *tK^UUU^* and *tQ^UUG^* could also partially rescue the defects in pathogenicity caused by the absence of *PoELP3* (Fig. 1-D and E). These results suggested that the defects caused by the deletion of *PoELP3* were at least partially caused by the inefficiency in protein biosynthesis.

### 3.2. PoElp3 is involved in the proper expression of protein

To further investigate how PoElp3 regulates fungal development, pathogenicity, and autophagy by affecting protein biosynthesis efficiency, we compared the protein abundance between Δ*Poelp3* strain and the Guy11 wild type strain through the tandem mass tags (TMT)-based quantitative proteomic analysis. Among the 5,165 proteins identified and quantified in our proteomic data, 386 proteins were significantly down-regulated [fold change (FC) > 1/1.3 (*p*-*value* < 0.05)], and 403 proteins were up-regulated [FC < 1.3 (*p*-*value* < 0.05)] (Fig. 2-A and B; Appendix D and E). The volcano plot result showed that most of the differentially accumulated proteins (DAPs) with a FC ranged from 1.3 to 2.0, and only a few of them with a FC >2.0 (Fig. 2-B). The clustering result showed that the DAPs could be divided into different groups according to their differential abundance between Guy11 and Δ*Poelp3* strain (Fig. 2-C).

**Fig. 2.**
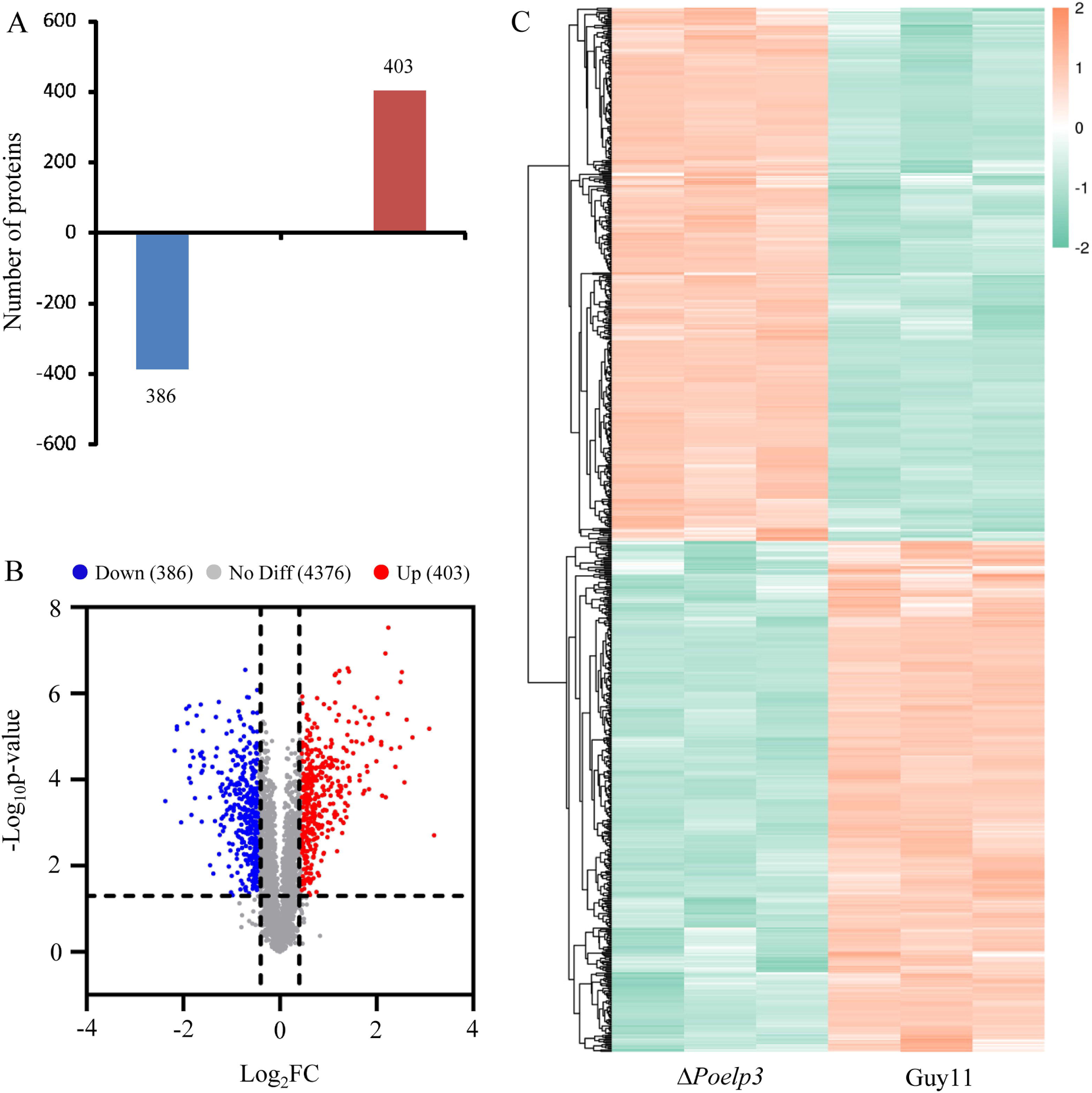
The differentially accumulated proteins (DAPs) between Δ*Poelp3* and Guy11 strains. A, quantitative identification of proteins upregulated and downregulated in Δ*Poelp3* strain though TMT. B, volcano plot showing the fold change of the differentially accumulated proteins. C, cluster analysis of the DAPs.

### 3.3. The DAPs involved in different pathways

To perform functional classification of the DAPs, we divided the DAPs into 4 Categories: Q1 with a FC<1/1.5, Q2 with a FC between 1/1.5 to 1/1.3, Q3 with a FC between 1.3 to 1.5, and Q4 with a FC>1.5 (Fig. 3-A). KEGG enrichment results showed that the Q1 DAPs were enriched in pathways such as Glyoxylate and dicarboxylate metabolism, biosynthesis of unsaturated fatty acids, various types of N-glcan biosynthesis, sulfur metabolism and sphingolipid metabolism. The Q2 DAPs were mainly enriched in steroid biosynthesis, tryptophan metabolism, and terpenoid backbone biosynthesis. While the Q3 and Q4 DAPs were mainly enriched in pathways such as aminoacyl-tRNA biosynthesis, biosynthesis, and metabolism of amino acids (Fig. 3-B). Protein domain enrichment showed that the Q1 DAPs were mainly enriched in pathways such as adenylysulphate kinase, fatty acid desaturase, and pyridine nucleotide-disulphide oxidoreductase. The Q2 DAPs were mainly enriched in pathways such as dynamin, flavodoxin, enoyl-(axyl carrier protein) reductase, and alpha amylase. While the Q3 and Q4 DAPs were mainly enriched in pathways such as tRNA synthetases, pyridoxamine 5’-phosphate oxidase, NADH oxidase and oxidoreductase NAD-binding domain (Fig. 3-C).

**Fig. 3.**
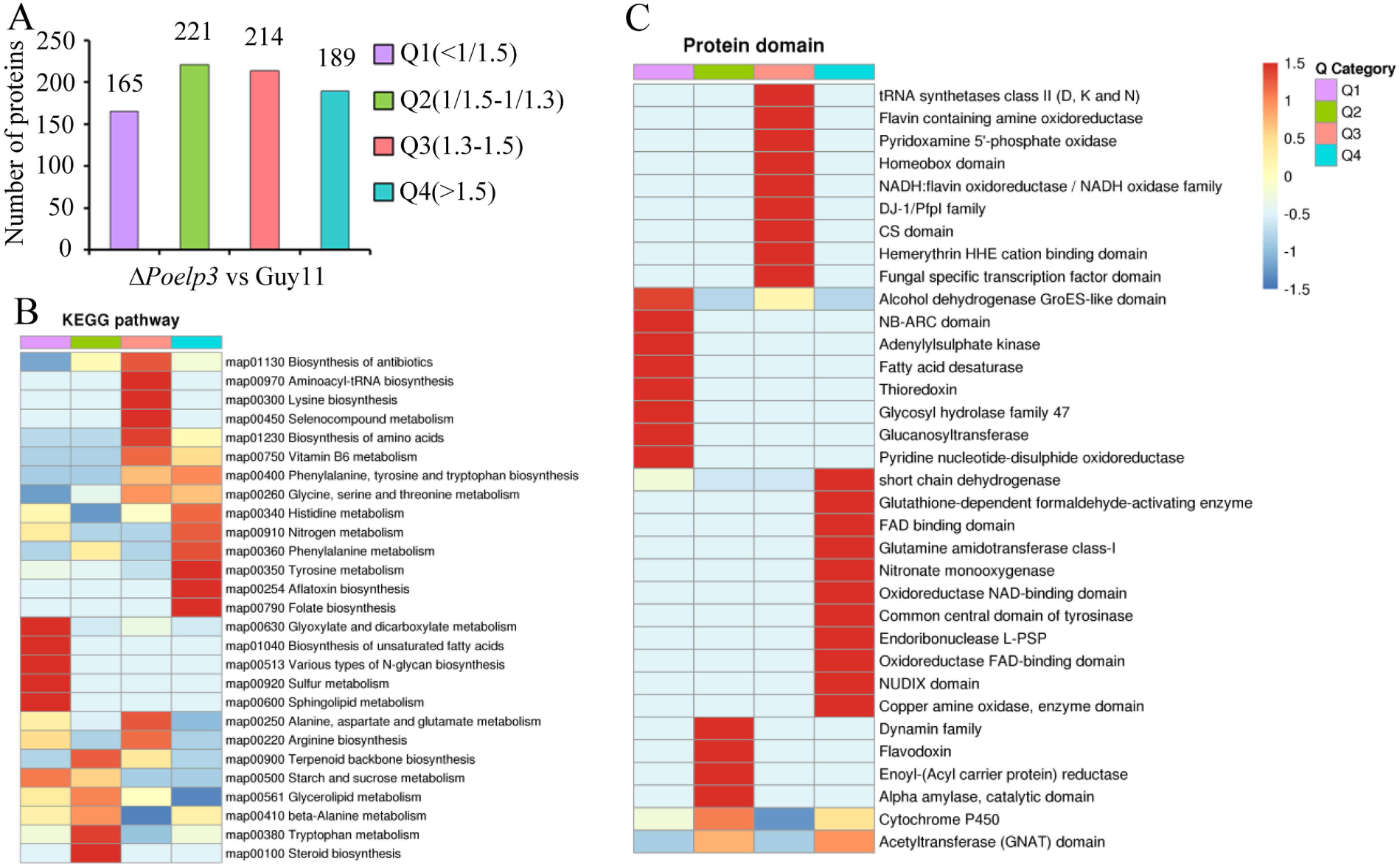
Functional classification of the DAPs. A, the four Q Categories of the DAPs. B, classification of DAPs according to the KEGG pathway. C, classification of DAPs according to protein domain.

### 3.4. PoElp3 regulated the accumulation of proteins independent of transcription

To further assess whether the DAPs were associated with the differential transcription level of their coding genes, we performed a comparative transcriptome analysis between the wild type and Δ*Poelp3* strains. We showed that only 535 genes were down-regulated, and 1,049 genes were up-regulated in Δ*Poelp3* compared with the wild type strain Guy11 (Fig. 4-A). We further investigated the transcription level of DAPs down-regulated in Δ*Poelp3* strain, and found that, among the 386 proteins down-regulated in Δ*Poelp3* strain, only 19 and 25 of them showing either higher or lower transcription level in the mutant strain (Fig. 4-B; Appendix F). While among the 403 DAPs up-regulated in Δ*Poelp3* strain, only 22 showing lower transcription level in the mutant strain, but 138 showing higher transcription level in the mutant strain (Appendix G), which could be affected indirectly. Taken together, these results suggested that the protein level of most of the DAPs in Δ*Poelp3* strain was independent of the transcription level.

**Fig. 4.**
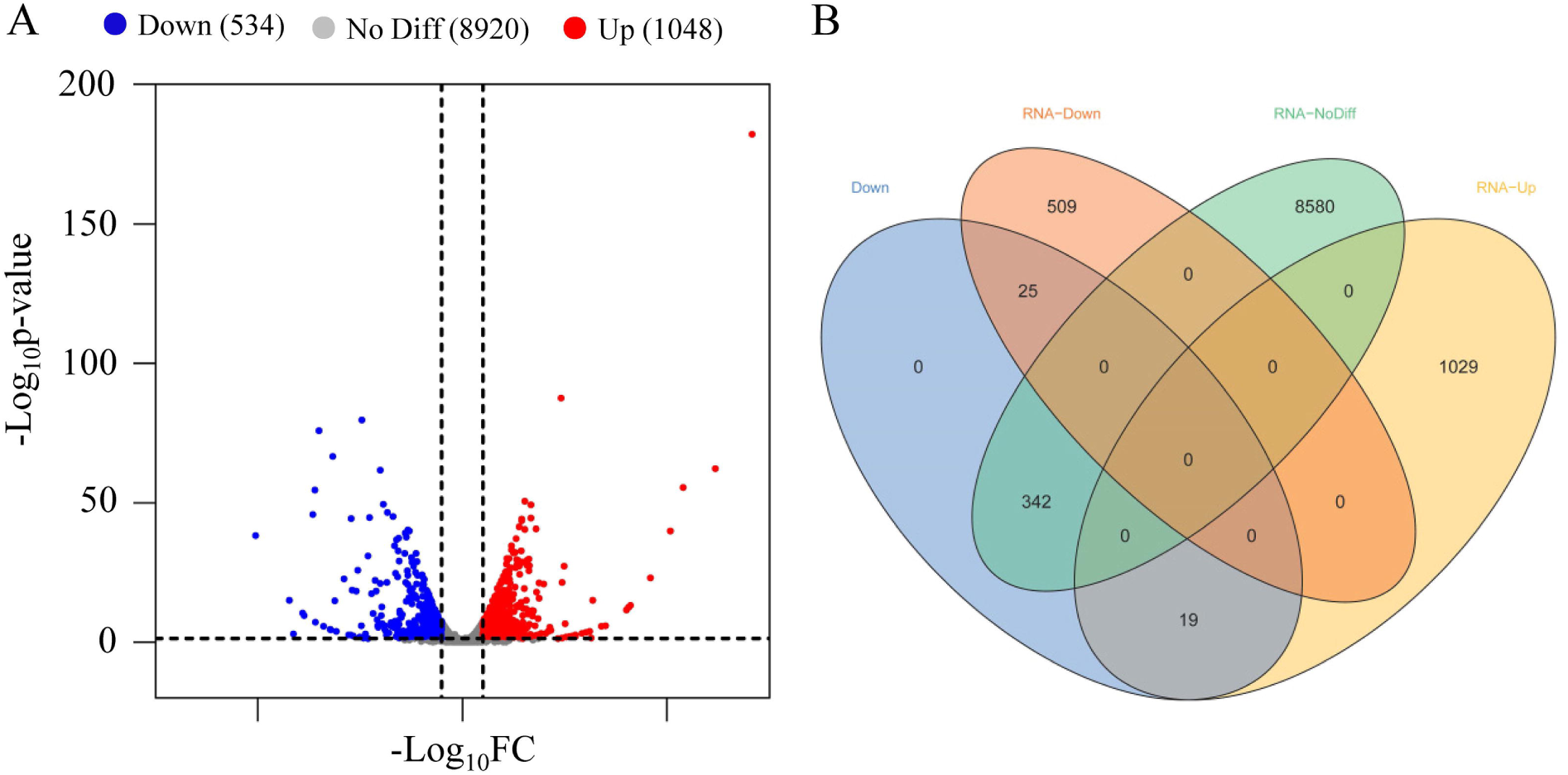
Comparative transcriptional analysis between wild type and Δ*Poelp3* strains. A, differentially expressed genes between wild type and Δ*Poelp3* strains. B, comparative analysis of the transcription level of DAPs down-regulated in Δ*Poelp3* strain.

### 3.5. Enrichment of Glutamine CAA-biased mRNA in the DAPs down-regulated in Δ*Poelp3* strain

To test whether there is a bias toward the use of NAA codons, we first surveyed the coding sequence of 70-15 genome (whole genome, WG), and found that 65.0%, 52.5% and 58.0% of the coding sequences showed an NAA:NAG ratio > 1 for Lysine, Glutamine, and Glutamic acid, respectively (Table 1). We then surveyed the coding sequence of DAPs down-regulated in Δ*Poelp3* strain. Since the DAPs down-regulated in Δ*Poelp3* strain were supposed to be caused by the absence of PoElp3, and are therefore referred to as Po<u>E</u>lp3 regulated <u>p</u>rotein candidates (Erps) hereafter. We found that 58.8.0%, 64.8% and 50.5.0% of the coding sequences showed an NAA:NAG ratio > 1 for Lysine, Glutamine, and Glutamic acid, respectively (Table 1; Appendix H). These results suggested an enrichment of Glutamine CAA-biased mRNA in the Erps compared with 70-15 genome.

**Table 1.**
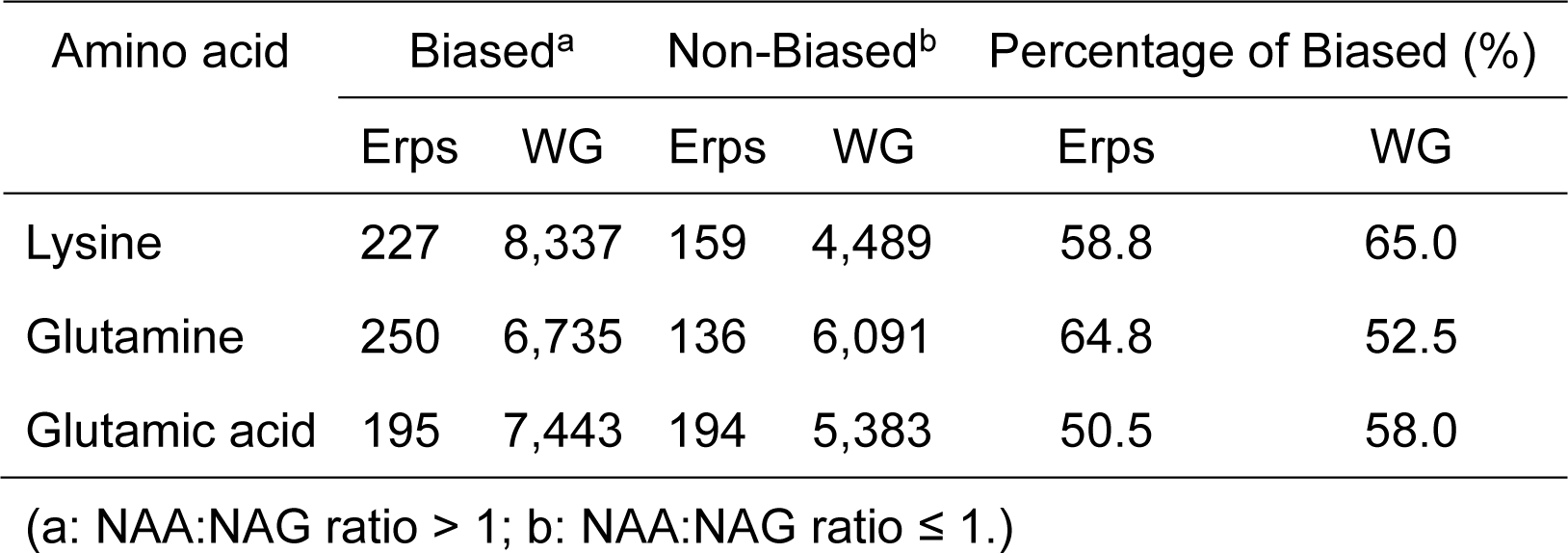
Codon usage of lysine, glutamine and glutamic acid in the Erps.

### 3.6. The Elp3-regulated proteins involved in asexual development, pathogenicity and autophagy

To assess whether any of the DAPs down-regulated in Δ*Poelp3* strain functions similarly to PoElp3, we searched in the PubMed database for publications associated with the 386 DAPs and found six DAPs involved in vegetative growth, conidiation, pathogenicity and/or autophagy. These DAPs presented a Fold Change (FC) ranging from 0.49∼0.76, and their transcription level in Δ*Poelp3* strain was no less than that in Wild type strain Guy11 (Table 2 and Appendix F). Among them, *ERP1* encodes a core protein of ESCRT-III subcomplex (Snf7) involved in asexual development, pathogenicity and autophagy (Cheng *et al*. 2018). *ERP2* encoding a Rho3 homolog is involved in appressorial development and pathogenicity (Zheng *et al*. 2007). *ERP3* encodes an ADP-ribosylation factor 6 protein (Arf6), which is involved in endocytosis and polarity during the asexual development (Zhu *et al*. 2016). *ERP4* encodes a tetrahydroxynaphthalene reductase (Buf1) and is required for melanin biosynthesis and full virulence (Zhu *et al*. 2021). *ERP5* encodes an amidophosphoribosyltransferase (Ade4), which catalyzes the conversion of 5-phosphoribosyl-1-pyrophosphate into 5-phosphoribosyl-1-amine in the de novo purine biosynthetic pathway and is required for conidiation and pathogenicity (Aron *et al*. 2022). *ERP6* encoding a glyoxylate aminotransferase 1 (Agt1) is required for mobilization and utilization of triglycerides during infection (Bhadauria *et al*. 2012a; Bhadauria *et al*. 2012b). Besides, we also found DAPs with their homologs in other fungal species associated with development or pathogenicity. For example, the MGG_03393 encodes a secondary metabolism regulator (Lae1) and is involved in asexual development and mycoparasitism of *Trichoderma atroviride* (Karimi Aghcheh *et al*. 2013). MGG_00413T0 encoding a putative β-1,3-glucanosyltransferase (Gel7) is required for the cell wall integrity under stress conditions (Zhao *et al*. 2014).

**Table 2.**
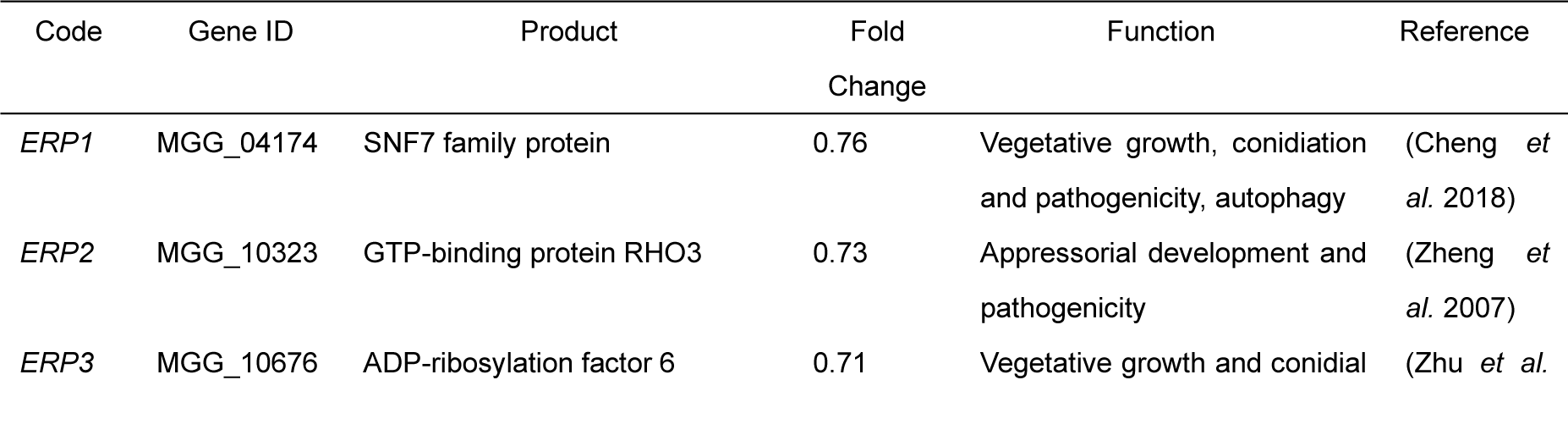
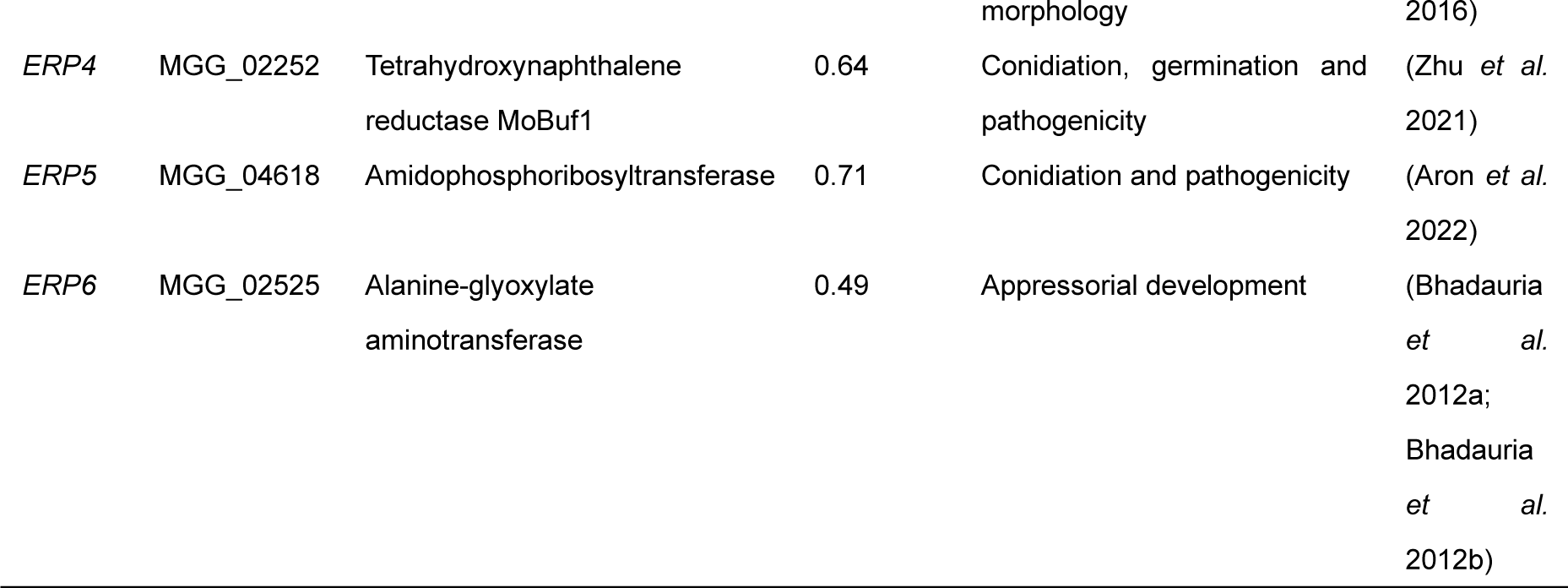
The *ERPs* functionally studied in *P. oryzae*.

### 3.7 The accumulation of PoErp9 was positively regulated by PoElp3

We further investigated the regulation of PoErp9, a sphingolipid C9-methyltransferase (MGG_14886), by PoElp3. To exclude the discrepancy caused by different genomic environment, the *PoERP9-GFP* fusion was fused *in situ* with the sulfonylurea-resistant allele of the *ILV2* gene (Yang and Naqvi 2014) in both the wild type Guy11 and Δ*Poelp3* strain (Fig 5-A). Immunobloting results showed that the abumdance of PoErp9-GFP in the Δ*Poelp3* strain was reduced to about half of that in wild type stain (Fig. 5-B; Appendix I and J), while the transcription level of *PoERP9* was not significantly reduced in the mutant strain (Fig. 5-C). These results suggested that PoElp3 is required for the translational efficience of PoErp9 independent of transcription.

**Fig. 5.**
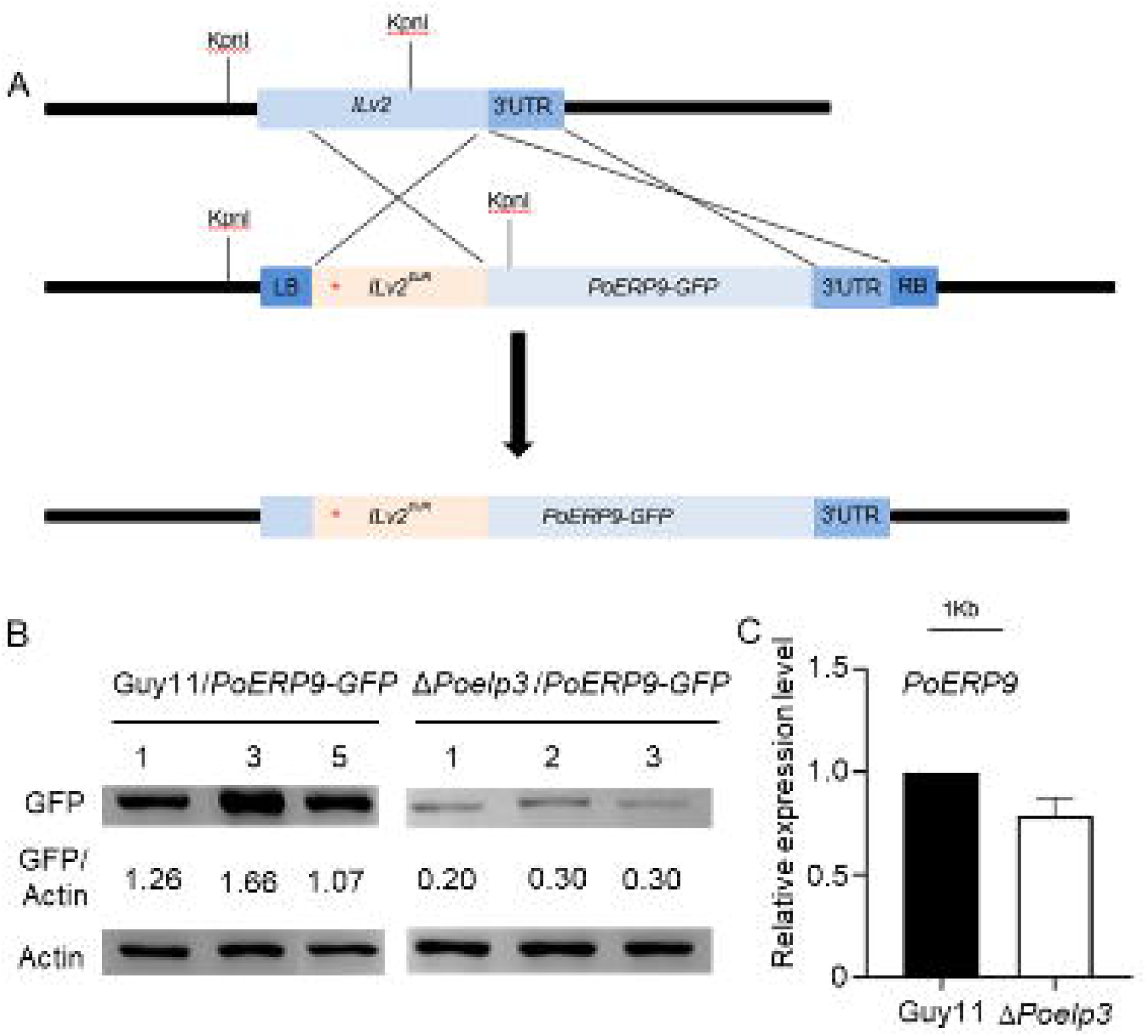
The abundance of PoErp9 was reduced in the Δ*Poelp3* strain. A, strategy for the *in situ* fusion of *PoERP9-GFP* into the sulfonylurea-resistant allele of the *ILV2* gene. B, the relative abundance of PoErp9-GFP [GFP/(GFP+GFP-PoAtg8)] in wild type (Guy11) and Δ*Poelp3* strains. The numbers indicated different individual transfromants. C, relative transcription level of *PoERP9* in wild type (Guy11) and Δ*Poelp3* strains based on qRT-PCR.

### 3.8 PoErp9 was required for the development and pathogenicity

To assess whether PoErp9 is functionally similar to PoElp3, we generated the Δ*Poerp9* strain using the split-marker strategy (Appendix C). The Δ*Poerp9* strain displayed reduced fungal growth and conidiation by compared with that of the *Ku80* parental strain and the complemented strain (Δ*Poerp9c*) (Fig. 6-A and C). The absence of *PoERP9* also caused defect in appressorial development and pathogenicity in the blast fungus (Fig. 6-D and F). The Δ*Poerp9* strain was less aggressive on the NPB leaves and generated mostly level 1 and level 2 lesions, while the *Ku80* parental strain and the complemented strain generated more lesions of level 3-5 (Fig. 6-E and F). These results suggested that PoErp9 is required for the fungal growth, appressorial development and the full virulence of the rice blast fungus.

### 3.9 PoErp9 act synergistically with the TOR-signaling pathway

To assess whether there is a interaction between PoErp9 and the Target-of-Rapamycin (TOR)-signaling, we performed rapamycin inhibition assays. Upon the treatment of rapamycin, the fungal growth of all the strains grown on either CM or MM media were reduced, and Δ*Poerp9*c strain displayed a higher inhibition rate by rapamycin compared with the parental strain *Ku80* and the complementation strain Δ*Poerp9c* (Fig. 7-A and B). We further detected the effect of rapamycin on the development of invasive hyphae (IHs) at the absence of *PoERP9*. In contrast to the parental strain *Ku80* and the complementation strain, the development of IH in Δ*Poerp9* strain was more severely inhibited by rapamycin (Fig. 7-C), and generated fewer type III and type IV IHs (Fig. 7-D). Taken together, these results suggested a role of PoErp9 in the interaction with the TOR-signaling during the vegetative and invasive growth of the rice blast fungus.

**Fig. 6.**
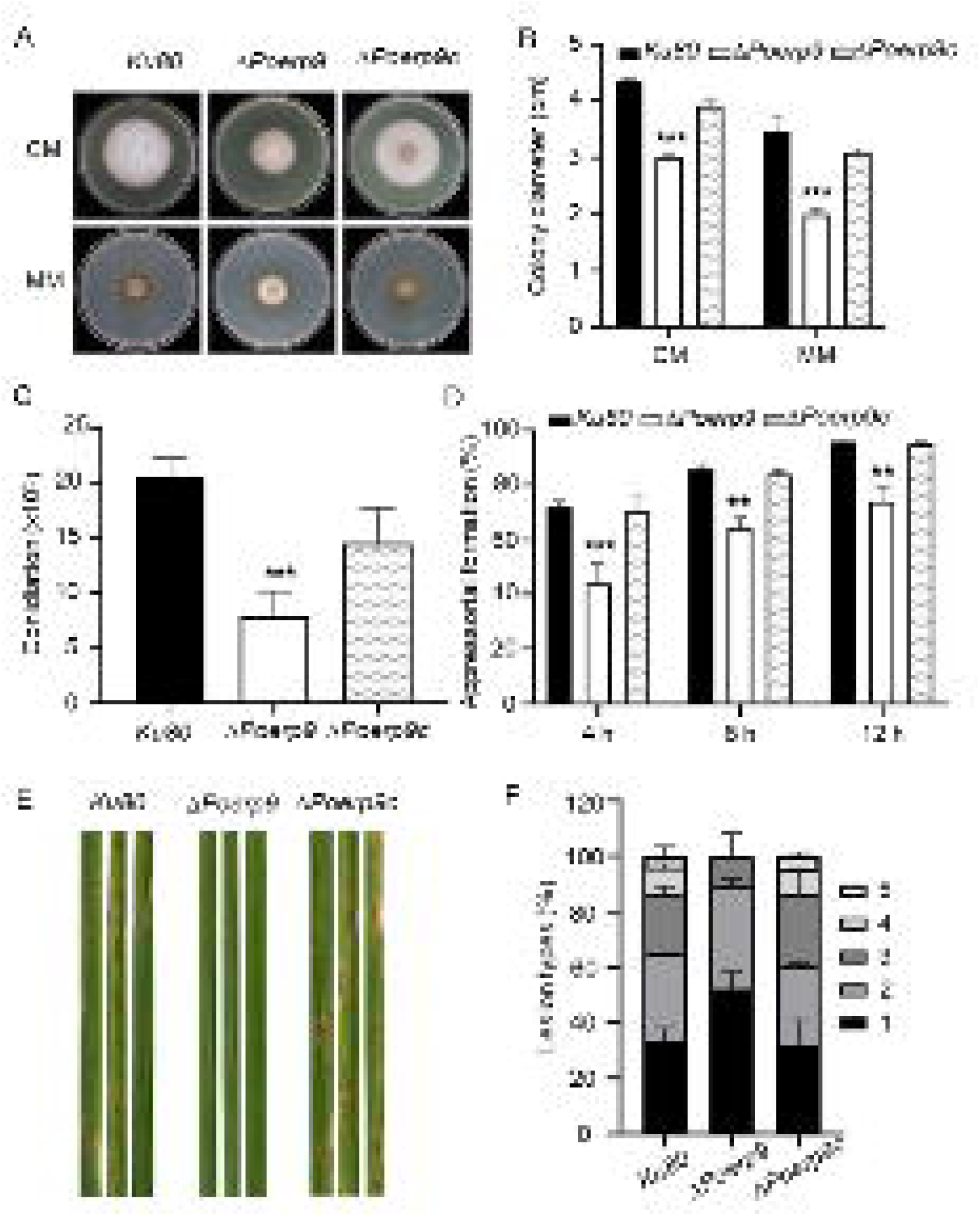
The Poerp9 strain was defective in asexual development and pathogenicity. A and B, the colony morphology (A) and colony diameter (B) of indicated strains on complete medium (CM) and minimal medium (MM) for 7 days. C, number of conidia harvested from a 7-cm rice bran plate at 3 days after induction of conidiation. D, appressorial development of Ku80, ΔPoerp9 and ΔPoerp9c was observed using microscope at 4, 8 and 12 h after induction. Data were shown as the mean±SE (n=3). The asterisks indicate statistically significant differences (**, P<0.01; ***, P<0.001). E, leaves of rice cultivar NPB were sprayed with conidial suspensions (5×104 conidia mL–1) of strains indicated. E, quantification of different lesion types. Data were shown as the mean±SE (n=3).

**Fig. 7.**
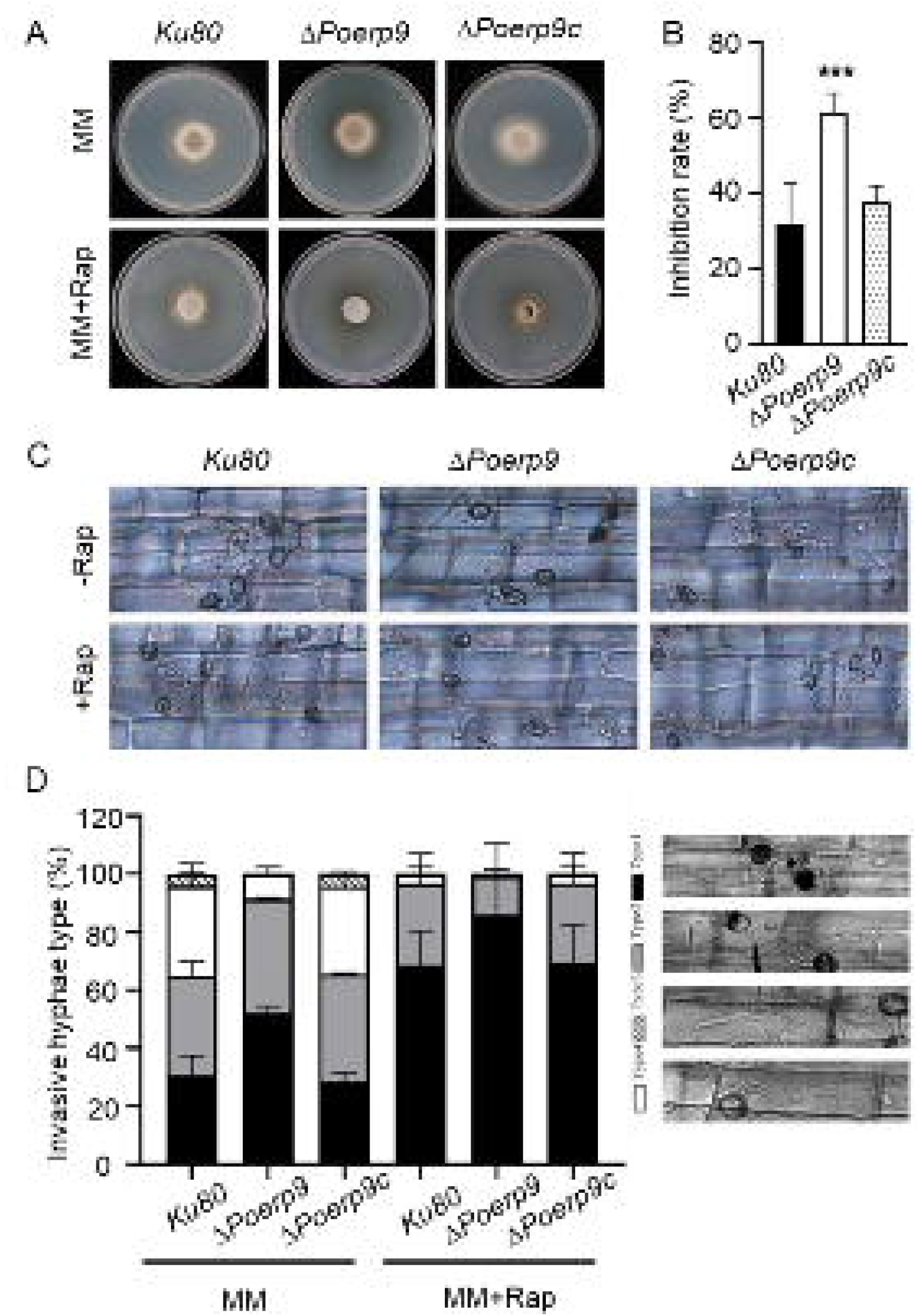
PoErp9 genetically interact with TOR-signaling to regulate vegetative and invasive growth of rice blast fungus. A and C, colony morphology (A) and inhibition rate of vegetative growth of indicated strains on minimal medium (MM) treated with rapamycin (Rap). C and D, morphology (C) and types (D) of invasive hyphae in *Ku80*, Δ*Poerp9* and Δ*Poerp9*c at 24 h post-inoculation (hpi). Data were shown as the mean±SE (n=3).

## 4. Discussion

The Elongator complex is conserved among different species and increasing evidence demonstrated that they function mainly through the modification of tRNA (Bauer *et al*. 2012; Candiracci *et al*. 2019; Dauden *et al*. 2019). In supporting this, overexpression of unmodified tRNA for glutamine, glutamic acid and lysine was efficient to restore the defects caused by the absence of Elongator in several studies (Esberg *et al*. 2006; Bauer *et al*. 2012; Zhang *et al*. 2022), probably by increasing the intracellular concentration of tRNAs to compensate for decreased binding affinity in the Elongator mutants (Esberg *et al*. 2006; Nedialkova and Leidel 2015; Tükenmez *et al*. 2015; Sokołowski *et al*. 2018). In this study, the phenotypic analysis showed that overexpression of *tK*(*UUU*) or *tQ*(*UUG*) could rescue fully the defects in fungal growth, and partially the defects in pathogenicity and response to rapamycin caused by the deletion of *PoELP3*, which indicated that PoElp3 may act through the tRNA-mediated protein integrity to regulate the fungal growth in rice blast fungus (Zhang *et al*. 2021).

Elp3 proteins had been proved to catalyze the modification of tRNA at wobbling U34 and consequently maintain the translation efficiency of proteins encoded by tK(UUU), tE(UUC), or tQ(UUG) (Bauer *et al*. 2012; Goffena *et al*. 2018). Previous studies suggested that NAA-biased codon usage is not always necessary for the Elongator regulated protein integrity via the tRNA modification. For example, the biased codon usage was exhibited in yeast (Bauer *et al*. 2012), but not in *A. fumigatus* (Zhang *et al*. 2022). In this study, we showed that the Erps displayed an enrichment of Glutamine CAA-biased mRNA, which suggested that different regulatory mechanisms might be applied in the Elp3 regulated translational efficiency. Moreover, the absence of NAA-biased cytoplasmic effectors on BIC as well as turnover of mRNA caused by the ribosomal pausing (Li *et al*. 2023) were not observed in Δ*Poelp3*. Possible explanations include that: 1) Elongator complex catalyzes the early step of mcm5s2 modification at U34 of tRNA, and 2) there are complementary pathways for the function of Elongator complex in the modification of U34.

The Elongator proteins could act by regulating the translational efficiency of target with important functions. For example, in fission yeast, the Elongator controls cell division by regulating the translational efficiency of the Cdr2 kinase (Bauer *et al*. 2012). Our functional assessment of the Erps showed that some of them were associated with asexual development, such as Erp1/Snf7, Erp2/Rho3, Erp3/ArF6, Erp4/Ade4, Erp5/Buf1, Erp6/Agt1, Lae1 and Gel7. Recent study in *A. fumigatus* showed that defects in development and pathogenicity in Δ*elp3* strain could be caused by the amino acid homeostasis imbalance (Zhang *et al*. 2022). We also found that many Erps were involved in the biosynthesis and metabolism of amino acids, suggesting a role of PoElp3 in development and pathogenicity by modulating amino acid homeostasis. Moreover, we also found Erps that may be involved in autophagy through different mechanisms. For example, Erp1/Snf7 acts in vesicle trafficking and autophagy (Cheng *et al*. 2018). Erp3/Arf6 is involved in autophagy homeostasis through the generation of phosphatidylinositol 4,5-bisphosphate (PIP(2)) and induction of phospholipase D (PLD) activity (Moreau *et al*. 2012). The absence of Erp4/Ade4 caused delayed glycogen mobilization and reduced turgor pressure in the appressoria (Aron *et al*. 2022), the representative cellular processes of autophagy during appressorial development in rice blast fungus (Deng *et al*. 2008; Kershaw and Talbot 2009). These results together suggested that PoElp3 is required for the development, pathogenicity and autophagy by regulating the translational efficiency of proteins involved in diverse pathways.

Both sphingolipid and ergosterol are important component of plasma membrane, and are involved in membrane trafficking (Hurst and Fratti 2020). Recent study in MoErg4 revealed a role in the proper localization of the *t*-soluble N-ethylmaleimide-sensitive factor attachment protein receptor (SNARE) protein MoSso1 and subsequently the secretion of cytoplasmic effector proteins in the rice blast fungus (Guo *et al*. 2023). Except for the involvement in autophagy, a common function in different organisms (Li *et al*. 2014; Liu *et al*. 2019), sphingolipid is also required for fungal growth (Ramamoorthy *et al*. 2009; Oura and Kajiwara 2010), and host - pathogen interaction (Ramamoorthy *et al*. 2009; Singh *et al*. 2012; Wang *et al*. 2021). In this study, functional classification revealed that some of the Erps were involved in sphingolipid metabolism. And detailed analysis suggested that PoElp3 is required for the translational efficiency of PoErp9, a sphingolipid C9-methyltransferase, and therefore maintains the proper metabolism of sphigolipid and the related cellular processes, including the asexual development, pathogenicity and TOR-related autophage homeostasis. However, how the PoErp9 act in these processed, particularly the pathogenicity, in the blast fungus remains to be explored.

In conclusion, our results demonstrated that PoElp3 was involved in the asexual development, pathogenicity, and autophagy, by regulating the tRNA-mediated protein integrity.

## Supporting information

Codon usage of lysine, glutamine and glutamic acid in the Erps

The ERPs functionally studied in P. oryzae

Integration of six graphs

## Acknowledgments

This work was supported by National Natural Science Foundation of China to H.Z (32172365) and Z.W (32272513), Central Guidance on Local Science and Technology Development Fund of Fujian Province (2022L3088), and the Innovative Research Funding of Fujian Agriculture and Forestry University, China, to H.Z (CXZX2020153D).

## Competing interests

The authors declare that they have no competing interests.

## Authors’ contributions

Z.W, G.L and H.Z conceived and designed the study. T.S, X.W and X.L performed data analysis. Y.H.L, Y.Y.L, J.H, H.B, J.H, Z.Z, and T.W performed the experiments. G.L, H.Z and Z.W wrote and revised the manuscript; GL, Z.W and D.Z contributed to improvement of the manuscript. All authors read and approved the final manuscript.

## Supporting information

**Appendix A.** Alignment of tRNAs from *Schizosaccharomyces pombe* (fission yeast) and *P. oryzae*.

**Appendix B.** Primers used in this study.

**Appendix C.** Generation of Δ*Poerp9* mutant.

**Appendix D.** Proteins up-regulated in Δ*Poelp3* strain.

**Appendix E.** Proteins down-regulated in Δ*Poelp3* strain.

**Appendix F.** The transcription level of DAPs down-regulated in Δ*Poelp3* strain. **Appendix G.** Comparative analysis of the transcription level of DAPs up-regulated in Δ*Poelp3* strain.

**Appendix H.** Codons used in Erps.

**Appendix I.** Southern bloting assays for the *ILV2*-specific PoERP9-GFP integration strains.

**Appendix J.** Ectopic expression of *PoERP9-GFP* could rescue the defect in fungal growth of Δ*Poerp9* strain.

## Notes

### Competing Interest Statement

The authors have declared no competing interest.

### Summary of Updates

Changed the order of authors. Added Figure 5p, Figure 6p, Figure 7p and Appendix.

