## Supplementary material for "Proteomic analysis revealed the function of PoElp3 in development, pathogenicity, and autophagy through the tRNA-mediated translation efficiency in the rice blast fungus": Codon usage of lysine, glutamine and glutamic acid in the Erps

Table 1 Codon usage of lysine, glutamine and glutamic acid in the Erps.

| Amino acid | Biased^a^ | | Non-Biased^b^ | | Percentage of Biased (%) | |
| --- | --- | --- | --- | --- | --- | --- |
|  | Erps | WG | Erps | WG | Erps | WG |
| Lysine | 227 | 8,337 | 159 | 4,489 | 58.8 | 65.0 |
| Glutamine | 250 | 6,735 | 136 | 6,091 | 64.8 | 52.5 |
| Glutamic acid | 195 | 7,443 | 194 | 5,383 | 50.5 | 58.0 |

(a: NAA:NAG ratio > 1; b: NAA:NAG ratio ≤ 1.)
