## Supplementary material for "Proteomic analysis revealed the function of PoElp3 in development, pathogenicity, and autophagy through the tRNA-mediated translation efficiency in the rice blast fungus": The ERPs functionally studied in P. oryzae

Table 2 The *ERPs* functionally studied in *P. oryzae*

| Code | Gene ID | Product | Fold Change | Function | Reference | |
| --- | --- | --- | --- | --- | --- | --- |
| *ERP1* | MGG_04174 | SNF7 family protein | 0.76 | Vegetative growth, conidiation and pathogenicity, autophagy | | (Cheng *et al.* 2018) |
| *ERP2* | MGG_10323 | GTP-binding protein RHO3 | 0.73 | Appressorial development and pathogenicity | | (Zheng *et al.* 2007) |
| *ERP3* | MGG_10676 | ADP-ribosylation factor 6 | 0.71 | Vegetative growth and conidial morphology | | (Zhu *et al.* 2016) |
| *ERP4* | MGG_02252 | Tetrahydroxynaphthalene reductase MoBuf1 | 0.64 | Conidiation, germination and pathogenicity | | (Zhu *et al.* 2021) |
| *ERP5* | MGG_04618 | Amidophosphoribosyltransferase | 0.71 | Conidiation and pathogenicity | | (Aron *et al.* 2022) |
| *ERP6* | MGG_02525 | Alanine-glyoxylate aminotransferase | 0.49 | Appressorial development | | (Bhadauria *et al.* 2012a; Bhadauria *et al.* 2012b) |
