## Supplementary material for "Proteomic analysis revealed the function of PoElp3 in development, pathogenicity, and autophagy through the tRNA-mediated translation efficiency in the rice blast fungus": Integration of six graphs

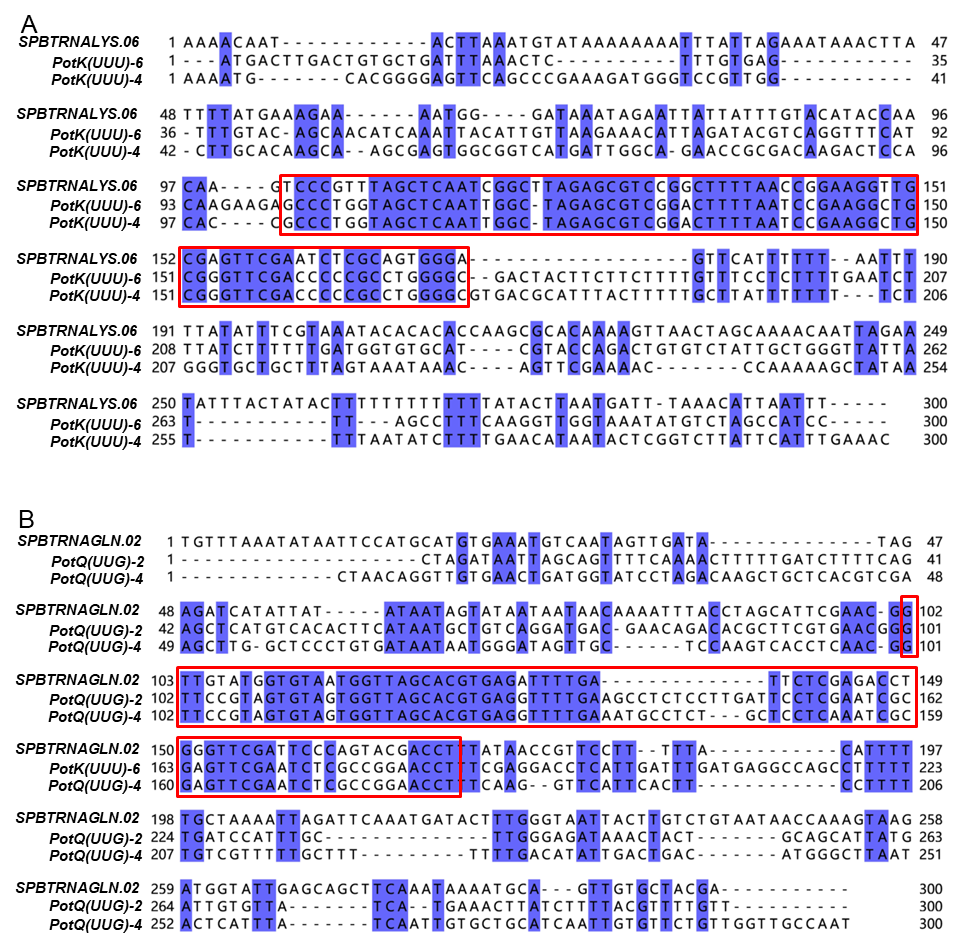

**Appendix A.** Alignment of tRNAs from *Schizosaccharomyces pombe* (fission yeast) and *P. oryzae*. (A). The alignment of tK(UUU) between *S. pombe* and *P. oryzae* (B). The alignment of tQ(UUG) between *S. pombe* and *P. oryzae*. The red rectangles indicate the coding sequences, and texts highlighted in blue show the conserved nucleotides.

**Appendix B.** Primers used in this study.

| Name | Sequence | Function |
| --- | --- | --- |
| MotQ-2F | GTCGACGGTATCGATAAGCTTAGCCAAGCATGGAAAATACCA | Amplification of tRNA for overexpression |
| MotQ-2R | TCAGTAACGTTAAGTGGATCCAATTGGGATGAGAGGGCGCGT |  |
| MotQ-4F | GTCGACGGTATCGATAAGCTTCTAAGTGCTCAGTTTCTCTGT |  |
| MotQ-4R | TCAGTAACGTTAAGTGGATCCAGCATATTGTTTTCGTGCCTG |  |
| MotK-4F | GTCGACGGTATCGATAAGCTTGAGTTCAGCCCGAAAGATGGG |  |
| MotK-4R | TCAGTAACGTTAAGTGGATCCTTCTAGGGTAAAGTTTTGCGT |  |
| MotK-6F | GTCGACGGTATCGATAAGCTTGACATGAAATCCTTTCAGTGTAG |  |
| MotK-6R | TCAGTAACGTTAAGTGGATCCCAATTTCCATACCTACCTCGC |  |
| *ERP9AF* | CGTTTGTGTTCTCGACCCTGA | Gene knockout |
| *ERP9AR* | TTGACCTCCACTAGCTCCAGCCAAGCCCACGGCGACACAAACGGGACT | Gene knockout |
| *ERP9BF* | GAATAGAGTAGATGCCGACCGCGGGTTCAAGAACCTCAACTGCGTCCA | Gene knockout |
| *ERP9BR* | GCTGATGATAGCACCCTCCGT | Gene knockout |
| *YG/F* | GATGTAGGAGGGCGTGGATATGTCCT | Gene knockout |
| *HY/R* | GTATTGACCGATTCCTTGCGGTCCGAA | Gene knockout |
| *HYG/F* | GGCTTGGCTGGAGCTAGTGGAGGTCAA | Gene knockout |
| *HYG/R* | AACCCGCGGTCGGCATCTACTCTATTC | Gene knockout |
| *ERP9tF* | GGAAGATTGGTGGTGGCACCA | Genotyping |
| *ERP9tR* | GATGAAGTCCTCGTACTGCCA | Genotyping |
| *ERP9UAF* | TTCATCGCTTCCGTCGCCTCA | Genotyping |
| *ERP9comF1* | agggaacaaaagctgggtaccAAAGAAGTTTGGTTGGGTGGT | Complementation |
| *ERP9comR1* | gcccttgctcaccataagcttTACTTTACGTCCCTCGCTCCA | Complementation |
| *ERP9qRT F1* | CACCGTTGGTGTTCACTACTC | qRT-PCR |
| *ERP9qRT R1* | CTGTACCACTTGTTGCCATACT | qRT-PCR |
| *ActinF* | CCATGTACCCTGGTCTTTCG | qRT-PCR |
| *ActinR* | TTCGAGATCCACATCTGCTG | qRT-PCR |
| *ERP9proslF2* | catgatgatgctcgagaattcAAAGAAGTTTGGTTGGGTGGT | PFGL1010-PoErp1-GFP |
| *ERP9ORFslR2* | CGCCCTTGCTCACGGTACCGCCTCCGCCGCCTCCGCCAGCCTTCTTGAGGTCAGGCAG | PFGL1010-PoErp1-GFP |
| *ILv2 F1* | ACCCGGCGTTGTTCTCGTCAC | Probe |
| *ILv2 R1* | ACAATACCTCCGCGTCCTTCAGC | Probe |

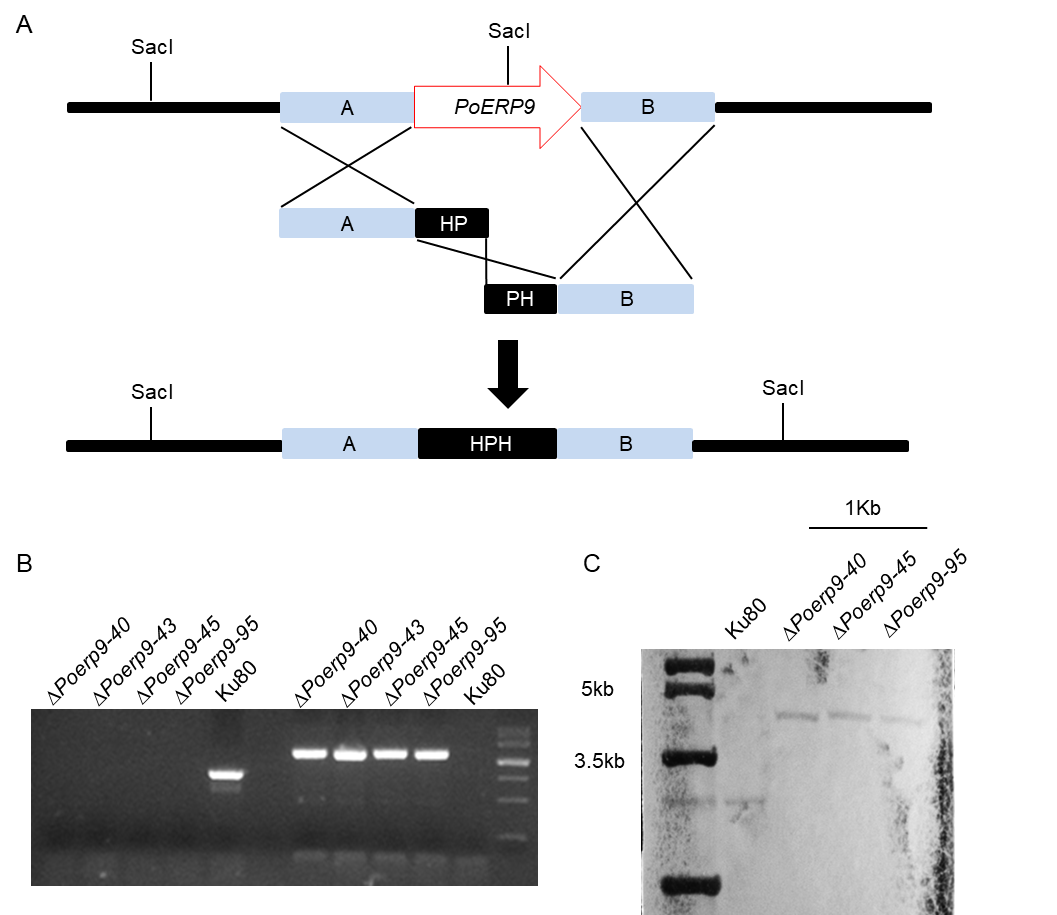

**Appendix C.** Generation of Δ*Poerp9* mutant. (A) Strategy for the mutagenesis of *PoERP9*. The A and B fragments of *PoERP9* were fused with N and C fragments of hygromycin phosphotransferase (HPH) genes respectively to form the 'A-H' and 'H-B' fragments that were used to replace the ORF of *PoERP9*. (B) Total genomic DNA samples (10 μg per sample) isolated from *Ku80*, *PoERP9* deletion mutants were digested with *Sac* I and subjected to Southern blot analysis using the 'A' fragment of *PoERP9* to generate the probe (Appendix B). A 4.8 kb target band was present each of the Δ*Poerp9* stains and a 2.8 kb target band was presented only in the *Ku80* parental strain.

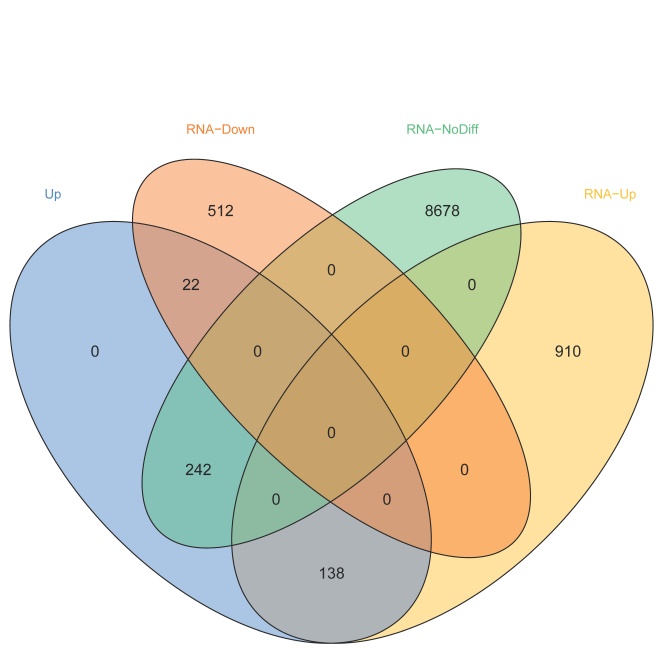

**Appendix G.** Comparative analysis of the transcription level of DAPs up-regulated in Δ*Poelp3* strain.

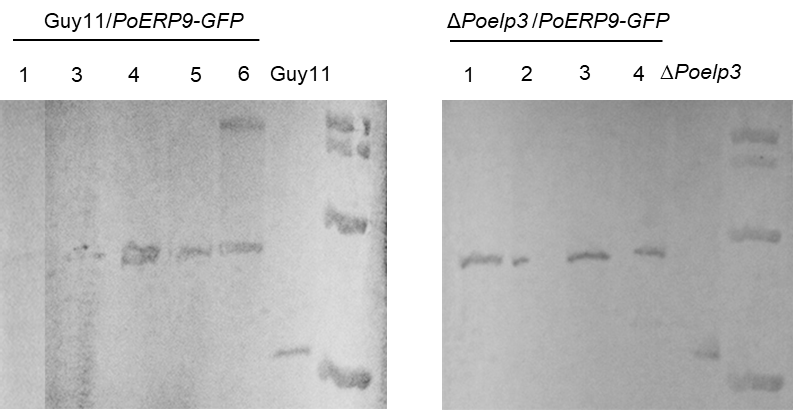

**Appendix I. Southern bloting assays for the *ILV2*-specific** *PoERP9-GFP* **integration strains.** Total genomic DNA samples (10 μg per sample) isolated from *Ku80*, *PoERP9* deletion mutant, and the indicated transformants were digested with *Kpn* I and subjected to Southern blot analysis using the 'A' fragment of *PoERP9* to generate the probe (Appendix B). A 2.2 kb target band was present in both the *Ku80* parental strain and Δ*Poerp9* strain, and a 3.4 kb target band was presented in the *PoERP9-GFP* *in situ* integration transformants.

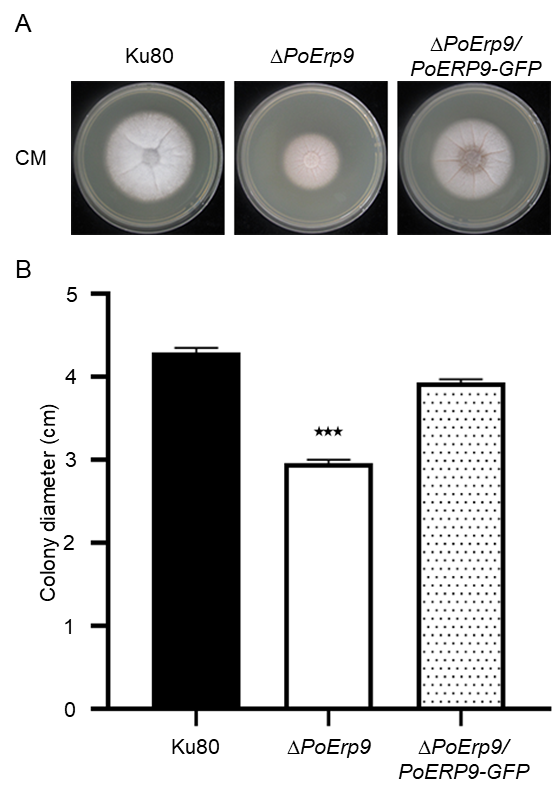

**Appendix J.** Ectopic expression of *PoERP9-GFP* could rescue the defect in fungal growth of Δ*Poerp9* strain.
